# Influence of STDP rule choice and network connectivity on polychronous groups and cell ensembles in spiking neural networks

**DOI:** 10.1101/2025.10.25.684525

**Authors:** Derek Arthur, Eric Albers, Masami Tatsuno

## Abstract

Spike-timing dependent plasticity plays an important role in how biological neural networks modify themselves with experience. However, the relationship between STDP and memory is not fully understood. Previously, an important advancement in understanding the relationship between spike-timing dependent plasticity (STDP) and memory was made through cortical simulations. The proposed memory items, polychronous groups (PGs), combined the network anatomy with precise spike-timing relationships between the connected cells of the network. However, there are some challenges with this previous work. It is unclear how different STDP rules would impact the PG results and if the PG results are complementary with purely spike-pattern defined memory items called cell ensembles (CEs). Lastly, it is unclear how these results are affected by changes in network connectivity. We address these challenges by comparing the PGs and CEs detected in spiking neural network simulations of cortical and hippocampal CA1 networks with two different STDP rule implementations. We show that the PG and CE results differ greatly for the two different STDP rules and for the cortical and CA1 networks. Our results show an important disconnect between anatomically defined and spike-pattern defined memories in spiking neural network simulations illustrating that care must be taken when drawing conclusions on the relationship between STDP and memory in simulation studies.

## 1. Introduction

The ability of biological neural networks to modify themselves with experience is central to understanding memory. This plasticity phenomenon and its connection to memory were first established when Donald Hebb proposed a general rule for synaptic modification, and how this rule led to groups of neurons coordinating their activity to represent memories: the cell assembly (Hebb, 1949). A crucial temporal component was added to Hebbian plasticity with the discovery of spike-timing dependent plasticity (STDP), where the change in the strength of a connection between neurons depends on the difference between the two cells’ spike timings (Bi & Poo, 1998).

One of the difficulties faced when performing in vivo recordings of neural activity is that we cannot measure the underlying anatomy simultaneously. This makes it difficult to understand the relationship between the changes in connection strengths and the patterns of cell activities present. To overcome this difficulty, simulations can be performed where the ground truth is known. Previously, an important advancement in understanding the relationship between network anatomy and memory was made through cortical simulations (Izhikevich, 2006). In his work, Izhikevich (2006) proposed a mechanism for the formation of memory called polychronization, where patterns of activity called polychronous groups (PGs) develop through precise spike-timing relationships among neurons sharing a common post- synaptic target. This initial work not only provided a connection between the network anatomy and memory but also demonstrated that the number of spike-timing patterns could far exceed the size of the network itself. However, there are three challenges in this work. The first is the type of STDP used in the simulation. They employed an early model of STDP known as additive-STDP (Song et al., 2000). However, it produces bimodally distributed connection strengths, with one end reaching zero strength and the other growing without limit. Therefore, the maximum strength that a single connection can achieve needs to be manually capped (Billings & Rossum, 2009; Rossum et al., 2000; Rubin et al., 2001; Song et al., 2000). The second is the use of PGs to represent memory. In electrophysiological experiments, PGs cannot be detected because they involve anatomical connections. Instead, researchers focus on detecting cell ensembles (CEs), statistically significant patterns of spiking activity among the recorded cells (Dechery & MacLean, 2017; Euston et al., 2007; Pérez-Ortega et al., 2021; Rolston et al., 2007; Sun et al., 2010; Vaz et al., 2023). Many detection methods for CEs have been developed (Mackevicius et al., 2019; Peyrache et al., 2009; Russo & Durstewitz, 2017; Watanabe et al., 2019), but it remains unclear how these CEs correspond to PGs. Finally, the original work investigated a simplified cortical network only. However, it remains unclear whether the findings from that network can translate to other types of networks.

We sought to answer these questions by (1) implementing another model of STDP, log-STDP (Gilson & Fukai, 2011), which produces lognormally distributed connection weights as has been observed in experiments (Song et al., 2005), (2) comparing the PGs to the CEs that are detected by an unsupervised method of detection (Russo & Durstewitz, 2017), and (3) comparing two network structures, one from the original cortical setup used by Izhikevich (2006) and the other corresponding to the hippocampal CA1 network.

## 2. Methods

### 2.1 Networks and Simulations

#### 2.1.1 Neuron Model

The networks are composed of Izhikevich model neurons (Izhikevich, 2003) with their dynamics determined by:

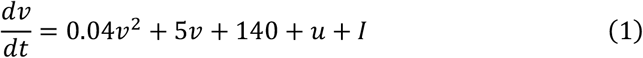

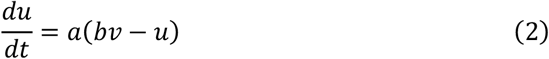

where *v* is the membrane potential, *u* is a membrane recovery variable representing the activation of K^+^ ionic currents and the inactivation of Na^+^ currents, *I* is the sum of synaptic inputs, and the dimensionless parameters *a,b,c*, and *d* determine the spiking activity. For all simulations the parameters for excitatory neurons are (*a, b*) = (0.02, 0.2) and (*c, d*) = (−65, 8) resulting in regular spiking dynamics. The parameters for inhibitory neurons are (*a, b*) = (0.1, 0.2) and (*c, d*) = (−65, 2) which give fast spiking type dynamics. A spike is recorded when *v* ≥ 30 *mV* after which *v* and *u* are reset: *v* ← *c* and *u* ← *u* + *d*.

#### 2.1.2 Cortex Network Structure

The cortex (CTX) network structure follows that of Izhikevich (2006). Each network is comprised of 800 excitatory and 200 inhibitory neurons, maintaining a 4:1 excitatory to inhibitory ratio observed in the cortex (Markram et al., 2004; Ren et al., 1992). The connectivity is determined by randomly selecting the post-synaptic targets of each neuron maintaining a connection probability of *p*_*ee*_ = 0.1 for excitatory-to-excitatory connections, *p*_*ei*_ = 0.1 for excitatory-to-inhibitory, *p*_*ie*_ = 0.1 for inhibitory-to-excitatory, and *p*_*ii*_ = 0 for inhibitory-to-inhibitory. All inhibitory connections are assigned a delay of 1 ms while the excitatory connections are assigned delay values ranging from 1 ms to 20 ms, with each delay chosen randomly from a uniform distribution. Lastly, in all cases, the weights are initialized so that all excitatory connections start with a weight of 6 mV while all inhibitory connections start with a value of −10 mV.

#### 2.1.2 CA1 Network Structure

The CA1 network connectivity is based on previous work (Azimi et al., 2021; Ramirez-Villegas et al., 2018). Each network was made up of 910 excitatory neurons and 90 inhibitory neurons to an approximate a 10:1 ratio. Each type of connection was given a unique connection probability. For excitatory-to-excitatory connections the probability was *p*_*ee*_ = 0.01, excitatory-to-inhibitory was *p*_*ei*_ = 0.30, inhibitory-to- excitatory *p*_*ie*_ = 0.25, and inhibitory-to-inhibitory *p*_*ii*_ = 0.25.

#### 2.1.3 STDP Rules

Two different STDP rules were used for this study. For additive STDP (add- STDP) (Song et al., 2000), the excitatory synaptic weights are updated according to

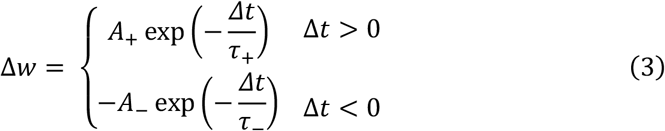

where Δ*t* = *t*_*post*_ − *t*_*pre*_ represents the time difference between the pre-synaptic spike arrival and post-synaptic spike emission. The decay constants for potentiation and depression are set to the same value *τ*_+_ = *τ*_−_ = 20 ms while the coefficients are given values of *A*_+_ = 0.1 and *A*_−_ = 0.12 to ensure the area under the depression curve is larger than the area under the potentiation curve. To prevent synapses from being potentiated indefinitely, a maximum value is set at 10 mV so that no connection weight can grow beyond this threshold.

The log-STDP rule (Gilson & Fukai, 2011) has the weights updated according to

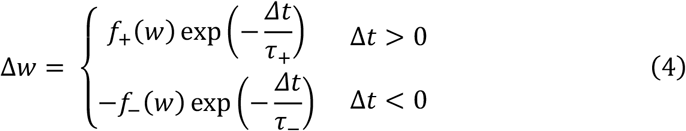

where the coefficients *f*_+_ and *f*_−_ become dependent on the current value *w* of the connection weight. These coefficients evolve via the equations:

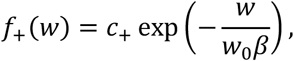

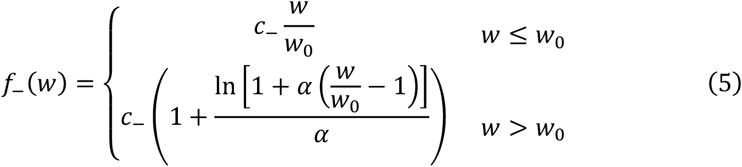

here *w*_0_ is a point chosen such that *f*_+_(*w*_0_) ≈ *f*_−_(*w*_0_), and the parameter *α* determines the log-like profile of *f*_−_ for weights larger than *w*_0_ (i.e., in the limit of *α* → 0, *f*_−_ reduces to the strictly linear equation 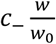). The dimensionless parameter *β* is chosen such that the potentiation coefficient decay does not allow a large portion of weights to become too strong. In such situations, the lognormal distribution is lost. For our simulations, a value of *β* = 2.33 maintains nearly all the synaptic weights at sub-spike threshold values. The dimensionless parameters *c*_+_ and *c*_−_ are set as 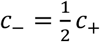 with *c*_+_ = 1.

#### 2.1.4 Simulations

For the simulations with no patterned input, a total of 20 independent simulations were performed for the cortex network as well as for the CA1 network. In both cases, 10 simulations used the log-STDP rule and the other 10 used the add-STDP rule. The connectivity is set randomly for each simulation, and the networks are driven by a 20 mV input to a randomly chosen neuron every millisecond. The synaptic weights are updated once every second. Each simulation is run for a total of 2 hours, where the first hour is to allow the network to settle into a state with a well-defined weight distribution and the second hour is to ensure analyses are performed on a 1-hour segment with no transient activities.

### 2.2 Polychronous Group Detection

Polychronous groups were detected using the same method as used by Izhikevich (2006). For a selected neuron, the detection algorithm searches for triplets of pre-synaptic neurons with connection weights of 9.5 mV or more, such that if all their spikes arrive simultaneously at the chosen neuron, a spike will be generated.

After finding all such triplets, a mini simulation is run for a preset duration *T*. The triplets are forced to spike at a time dictated by the assigned connection delay, after which all spontaneous activity caused by these activations is recorded until time *T* is reached. The algorithm will discard a result if there is no spontaneous activity beyond the one neuron that is forced to fire by stimulation of its three pre-synaptic connections. The maximum time for the mini simulation *T* was set to 150 ms. Example PGs are shown in Figure 2A. Lastly, the algorithm was run using the weight matrix corresponding to the 1^st^, 900^th^, 1800^th^, 2700^th^ and 3600^th^ second of each hour of each simulation.

### 2.3 Cell Ensemble Detections

The method used for detecting cell ensembles in the simulation spike data is one proposed by Russo and Durstewitz (2017). The method detects cell ensembles in an unsupervised manner for arbitrarily chosen time lags between spikes. The spike matrices are binned according to predetermined values and significance is tested for a range of lags preset by the user. The bins chosen for the simulations were 3, 5, 10, 15, 25, 30, and 50 ms. This range covers cases of precise spike timing sequences for the low bin values and imprecise spike sequences or groups of neurons with correlated firing rates at the larger bin values. The maximum lag for all bins was chosen as 10 bins; that is, the algorithm considers lags ranging from −10 to 10 bins from the reference spike train. This detection method comes with a large computational cost for large numbers of neurons. So, for every simulation we composed subsets of the spiking data comprised of the spiking activity of 30 randomly sampled neurons. We did this 10 times and ran the detection algorithm on each of these 10 subsets. Example CEs are shown in Figure 3A.

### 2.4 Determination of CE and PG lifetimes

PG lifetimes are directly related to the lifetime of connections with weights above 9.5 mV. Since the detection algorithm requires the connection weights between the three anchor neurons and the common post-synaptic target to be above this value, as soon as one of those three connections drops below 9.5 mV the PG ceases to exist. This means there is a 1-to-1 correspondence between PG lifetimes and the lifetimes of strong (>9.5 mV) connections in the simulation. CE lifetimes were calculated as the time difference between the first detected activation and last detected activation of a given CE.

### 2.5 Cross-correlation analysis

To quantify a relationship between the CEs and the connection weights in each network, we computed the cross-correlation between the number of active CEs each second and one of two other quantities. The first quantity is the total amount of excitatory weight *W* for each second, calculated as

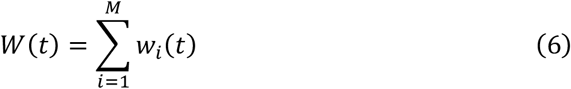

where *w*_*i*_(*t*) is the weight of the excitatory connection *w*_*i*_ at time *t* and *M* is the number of excitatory connections in the network. The second quantity cross-correlated with the number of active CEs each second is the number of excitatory connections each second with weights between specified 1 mV weight ranges. That is, the quantity 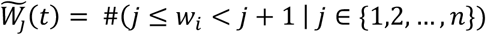 where *n* is the largest connection weight achieved in the network.

## 3. Results

Raster plots of typical network activity are shown in Figure 1A. After 1 hour of CTX simulations, the add-STDP networks exhibited dense background spiking activity with periodic bursts of highly synchronized spiking of 30-40 Hz; on the other hand, the log-STDP networks showed sparser background activity with periodic bursts of high firing rate activity. The spiking within these bursts was approximately 20-30 Hz in frequency and the burst occurrence rate ranged from 2-5 Hz. After 1 hour of CA1 simulations, both add-STDP and log-STDP displayed random spiking activity with no periods of increased firing rate or synchronization. The overall shapes of the distributions for excitatory connection weights were stable. For both the CTX and CA1 simulations with log-STDP, the weights were distributed lognormally with a peak at approximately 4 mV and no weight exceeding 24 mV (Fig. 1B). The weight distributions for the CTX simulations with add-STDP were bimodal with most weights pushed to the strongest value of 10 mV. In the CA1 simulations, nearly all excitatory weights were pushed to this maximum value.

**Figure 1.**
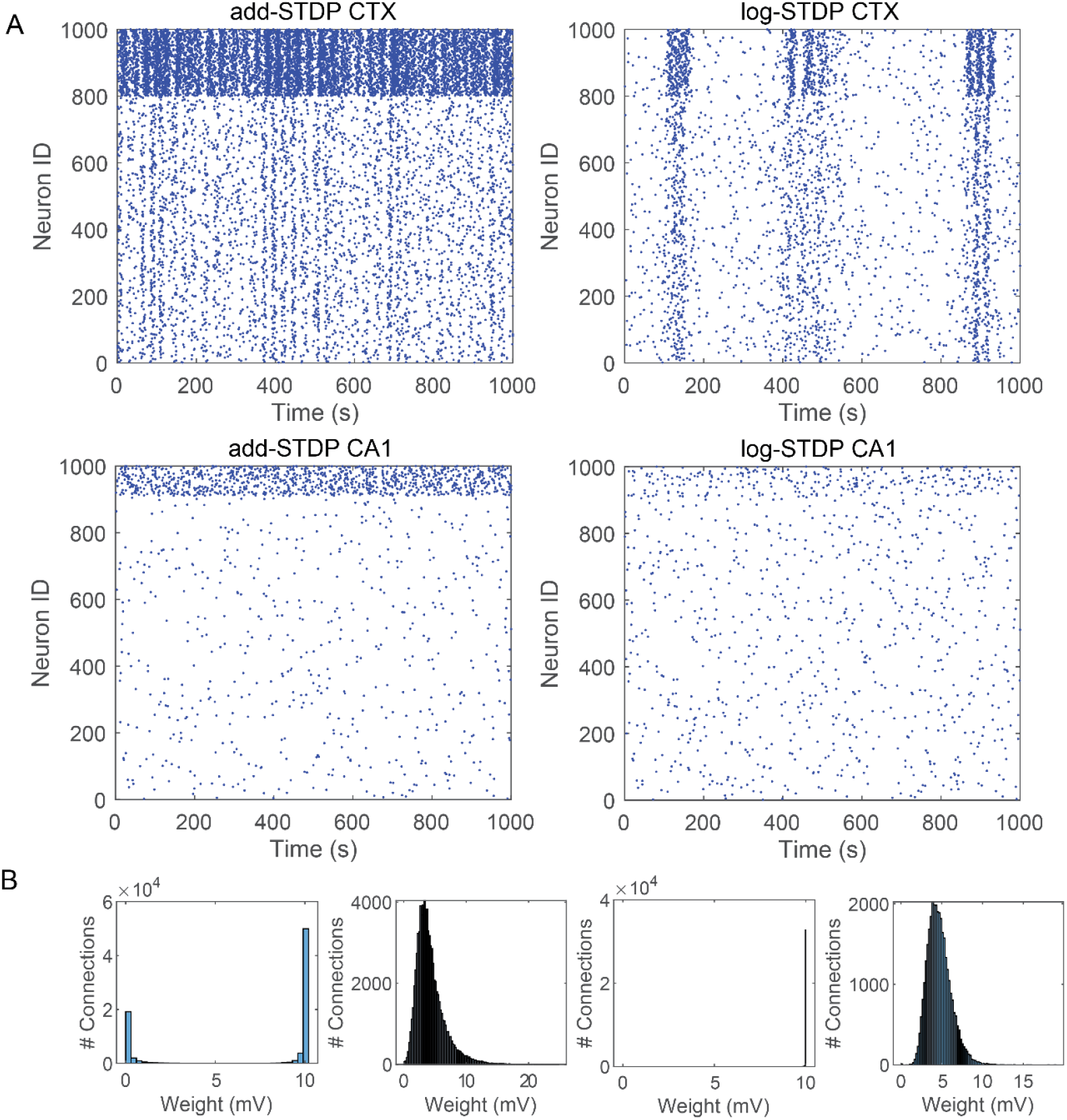
**A.** Raster plots over 1 second of spiking activity after 1 hour of simulation time. For the CTX simulations, the first 800 units are excitatory neurons, and the first 910 for the CA1 simulations. **B**. Example distributions of excitatory connection weights after 1 hour of simulation time.

**Figure 2.**
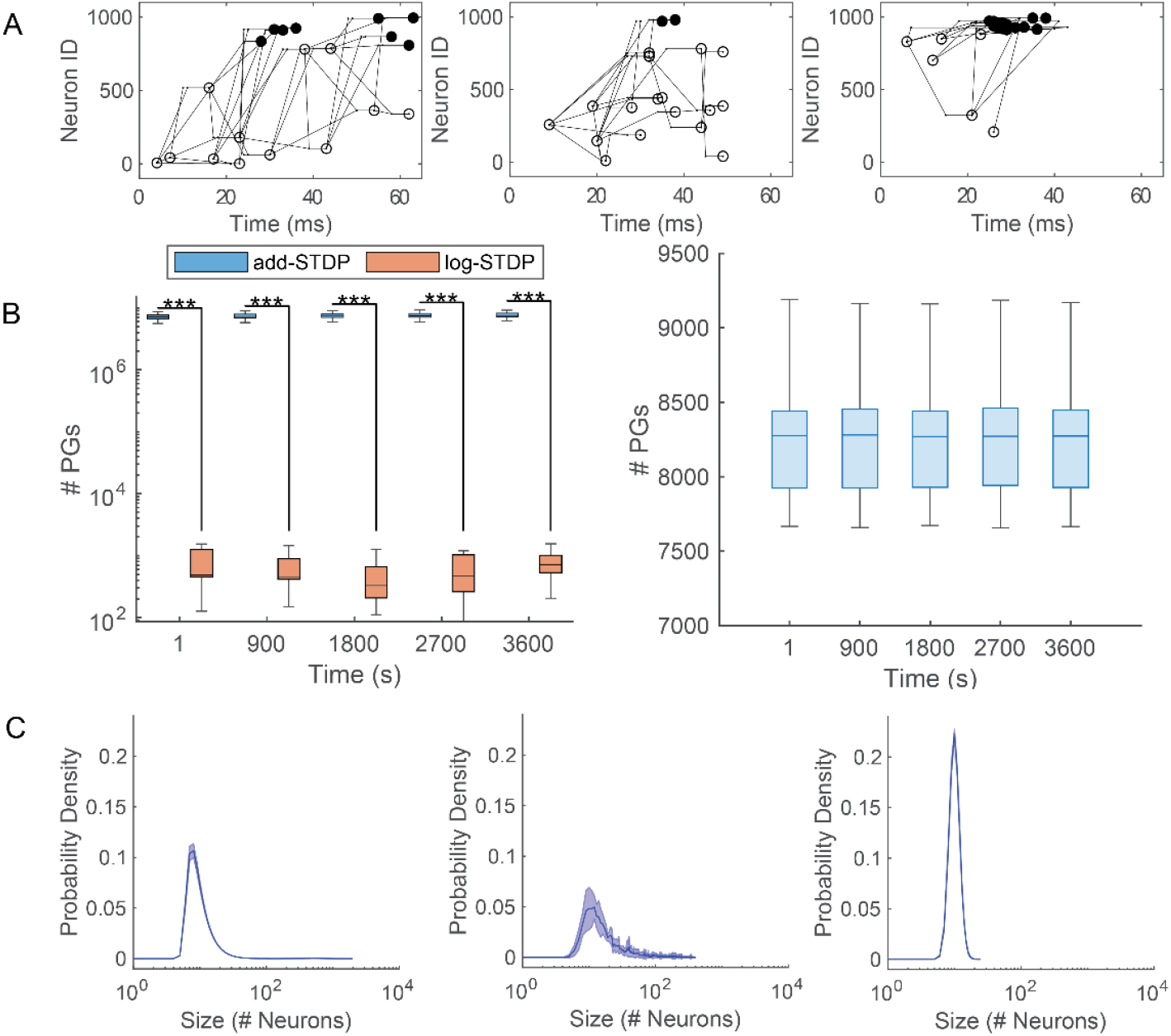
**A.** Example PGs for add-STDP in cortex (left), log-STDP in cortex (center), and add-STDP in CA1 (right). The first 800 neurons are excitatory and the last 200 are inhibitory. Open and closed circles represent excitatory and inhibitory neurons, respectively. **B**. Number of PGs detected at 15-minute intervals in cortex simulations (left) and in CA1 simulation with add-STDP (right). Box plots show the median and interquartile range (IQR) of PG counts, upper and lower whiskers represent the maximum and minimum PG counts, respectively. **C**. PG size distributions averaged over all simulations at t=3600 s for add-STDP CTX (left), log-STDP CTX (middle), and add-STDP CA1(right). Mean is shown by the solid line, and the shaded area represent 1 SD. Statistical analysis by Wilcoxon rank sum test, ***p<0.001.

**Figure 3.**
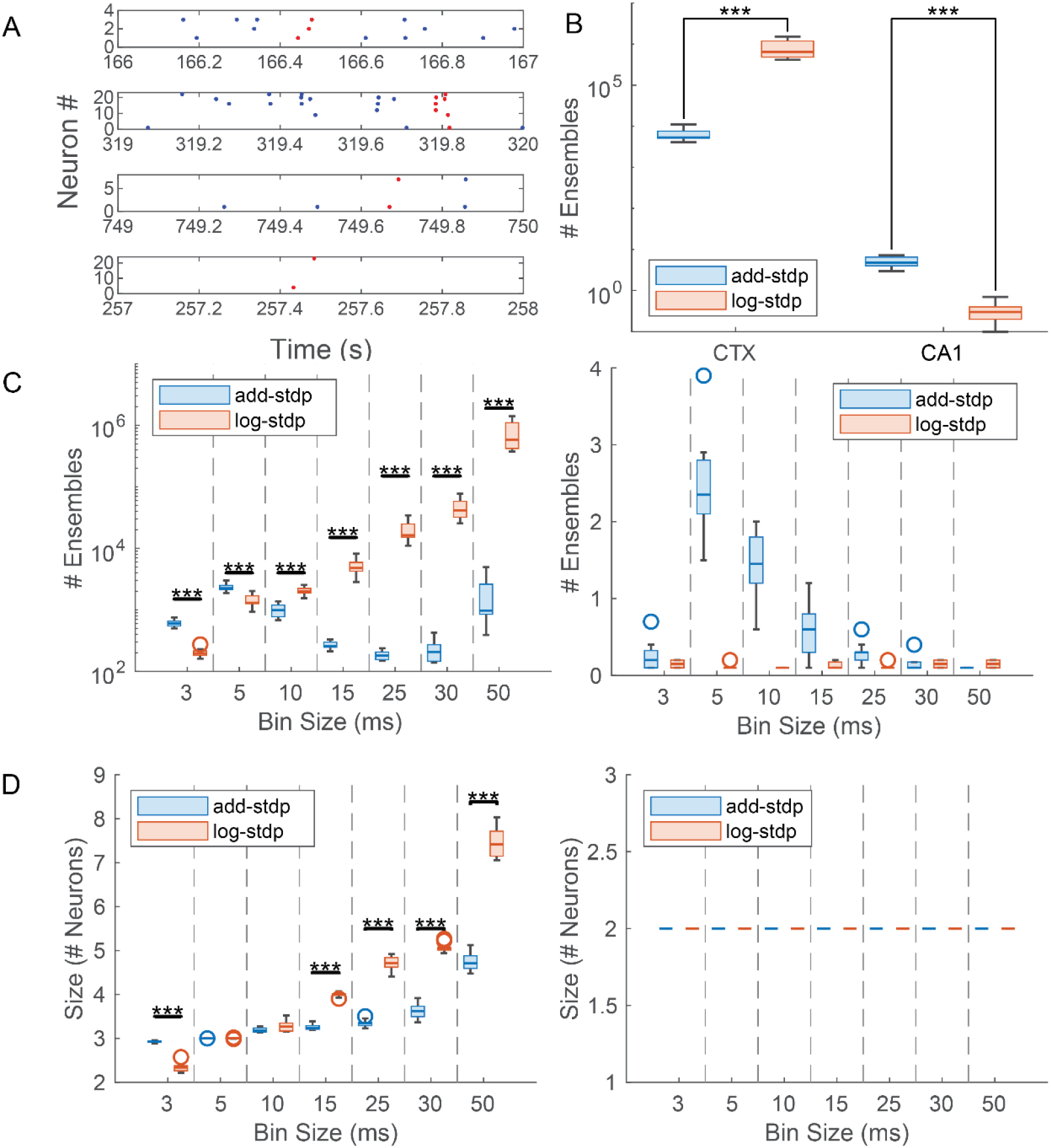
**A.** Example CEs. Red dots represent spikes constituting CE activation. **B**. Total number of CEs detected across 10 simulations **C**. Total CEs per bin for CTX simulations (left), and for CA1 simulations (right) **D**. Size of CEs per bin for CTX (left) and CA1 (right). Boxes show median and IQR, whiskers show non-outlier maximum and minimum values. The threshold for outlier detection was set to 1.5*IQR. Statistical analysis by Wilcoxon’s rank sum test, *p<0.05, **p<0.01, ***p<0.001.

### 3.1 More PGs detected in add-STDP simulations than log-STDP simulations

For the PG detection, the algorithm used the connection weights of 9.5 mV or more (see Methods). With a larger number of connections above this threshold with add-STDP, the simulation produced many more PGs than the log-STDP simulations. To investigate the stability of detection, we ran the detection algorithm at 5 different points in each simulation, spaced 15 minutes apart. In the CTX networks, the add- STDP simulations produced, on average, 10^4^ more PGs than the log-STDP simulations. For the CA1 networks, only the add-STDP simulations produced PGs (Fig. 2B). Next, we compared the sizes (number of member neurons) of the detected PGs. In the two CTX simulations and the CA1 simulation with add-STDP, the peak in the distributions of PG sizes were all in the range of 8-12 neurons. However, the maximum PG size differed considerably: approximately 2000 neurons for the add- STDP CTX simulation, 400-474 neurons for the log-STDP CTX, and 25 neurons or lower for the CA1 add-STDP. Example distributions of PG sizes averaged across all 10 simulations at a single time point are shown in Fig. 2C.

### 3.2 More CEs detected in log-STDP CTX simulations

If CEs and PGs are closely related, we expect to see higher CE detections in add-STDP than in log-STDP for the CTX simulations. However, we found the opposite (Fig. 3B). For each simulation we ran the CE detection algorithm on ten subsets of the spiking data. Each subset was composed of the spiking data for 30 excitatory neurons chosen randomly. The total number of CEs detected was averaged across these subsets for each simulation resulting in a distribution of 10 values for the total number of CEs detected. Comparing these resulting distribution of averages showed a significant difference (z = −3.7418, p = 1.8267×10^−4^; rank sum test) between the number of CEs for the log-STDP simulations (m = 6.47×10^5^, IQR 4.83×10^5^ – 1.19×10^6^) and the add-STDP simulations (m = 5323, IQR 5120-7594). For the CA1 simulations, the average number of detected CEs was significantly higher for add- STDP (m = 4.8, IQR 4-6.5) than log-STDP (m = 0.3, IQR 0.2-0.4), exhibiting an opposite trend to the CTX simulations (z = 3.7574, p = 1.7168×10^−4^; rank sum test). Making bin-wise comparisons of the number of detected CEs showed a difference in the preferred bin size for each rule (Fig. 3C). The add-STDP simulations for both the CTX and CA1 had the highest number of detected CEs occurring in the 5 ms bin, whereas the log-STDP cortex simulations showed the highest number of detected CEs in the 50 ms bin. The log-STDP CA1 simulation had too few CEs detected to determine a preferred bin size. Bin-wise comparisons of mean CE sizes were also different from the PG results (Fig. 3D). In the CTX simulations, the CE sizes ranged from 2 to 5 neurons for add-STDP and 2-8 neurons for log-STDP. For the CA1 simulations, all CE sizes were 2 neurons regardless of the STDP rule.

### 3.3 CEs are stable in all networks, PGs are unstable in log-STDP CTX simulations

We next investigated the stability of the detected CEs and PGs by checking their lifetimes. Distributions of CE lifetimes are shown in Figure 3A. Many of the detected CEs have a lifetime greater than 2000 seconds for both STDP rules and both networks. For the PG lifetimes, we assessed the lifetime of weights in the network that were larger than 9.5 mV, corresponding to the threshold in the PG detection algorithm (see Methods). The detection algorithm requires the connection weights between the common post-synaptic target and its three anchor neurons to be a value larger than 9.5 mV. If any of these three connections drops below 9.5 mV in value, then the PG cannot be detected and, therefore, no longer exists. Thus, PG lifetimes are directly related to the lifetime of connections with values above 9.5 mV in the networks. Distributions of these lifetimes averaged over all simulations are shown in Figures 4B & 4C. For the log-STDP cortex simulations the distribution peaks at 28 seconds, with the maximum lifetime being 1853 seconds. For the add-STDP CTX simulations, the strong weights were stable for 3600 seconds, which is the duration of the analyzed part of the simulations. In the add-STDP CA1 simulations, the strong connections were very stable with all having lifetimes of 3600 seconds. The strong connections in the log-STDP CA1 simulations were also very stable with all connections having a lifetime greater than 3956 seconds.

**Figure 4.**
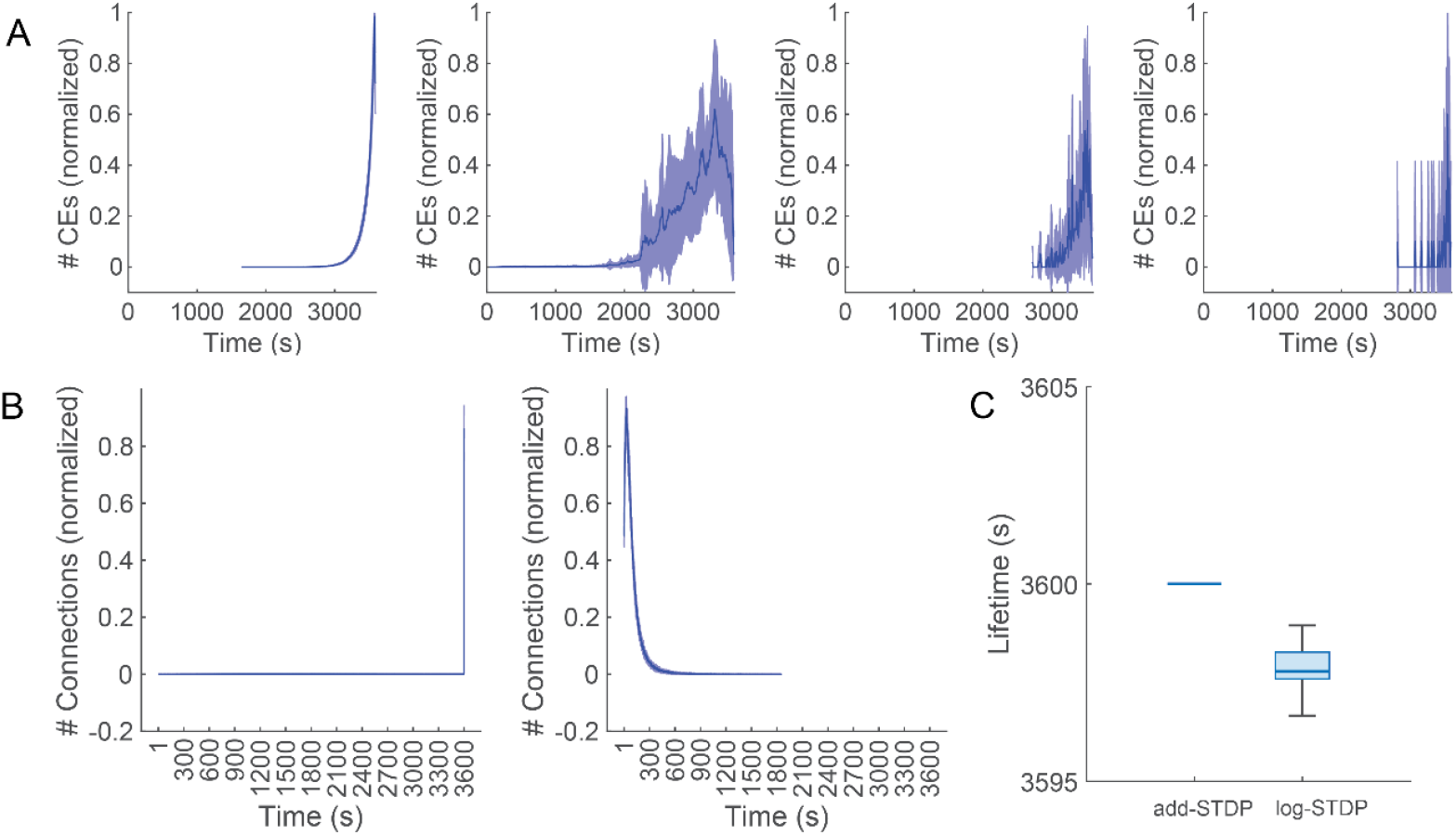
**A.** Distributions of CE lifetimes for add-STDP CTX (left), log-STDP CTX (middle-left), add-STDP CA1 (middle-right), and log-STDP CA1 (right) simulations. Mean is shown by the solid line, and the shaded area shows 1 SD. **B**. Distributions of strong (>9.5 mV) connection weight lifetimes in add-STDP CTX (left) and log-STDP CTX (right) simulations. Number of connections were max-min normalized to the range 0-1 from 0-17975 (add-STDP) and from 0-450 (log-STDP). Mean is shown by the solid line, and the shaded area shows 1 SD. **C**. Lifetimes of strong (>9.5 mV) weights in CA1 simulations. Boxes show median and IQR, whiskers show maximum and minimum lifetimes.

### 3.4 Amount of excitation anti-correlated with active CEs in log-STDP CTX simulations, weak correlations for add-STDP CTX

Finally, we investigated the temporal relationship between the dynamics of connection weights and those of CEs. If the weights provide a stage for CEs to be activated, the changes in weights are expected to precede the changes in CEs. Due to the low number of CEs detected in the CA1 simulations this analysis was only performed for the CTX simulations. Example time courses of the total amount of excitatory weight and the number of active CEs (ens-MUA) are plotted in Fig. 5A. In these representative examples, there was no clear correlation for add-STDP (Fig. 5A, left), but the two signals were anti-correlated for log-STDP (Fig. 5A, right). To quantify this observation, we extracted the lags corresponding to the largest peaks and the z-scored cross-correlation values from all simulations. For the CTX, the add-STDP simulations exhibited both positive and negative lags (Fig. 5B, left) and relatively small peak correlation values between 0 and −0.4 (Fig. 5B, right, blue bar). In contrast, the log-STDP simulations showed consistently negative lags of 12-16 seconds (Fig. 5B, middle) and relatively large peak correlation values around −0.8 (Fig. 5B, right, red bar). The negative lags and negative peaks indicate that an increase in the number of active CEs preceded a decrease in the total amount of excitatory weights. In other words, changes in CEs precede changes in weights, contrary to our prediction.

**Figure 5.**
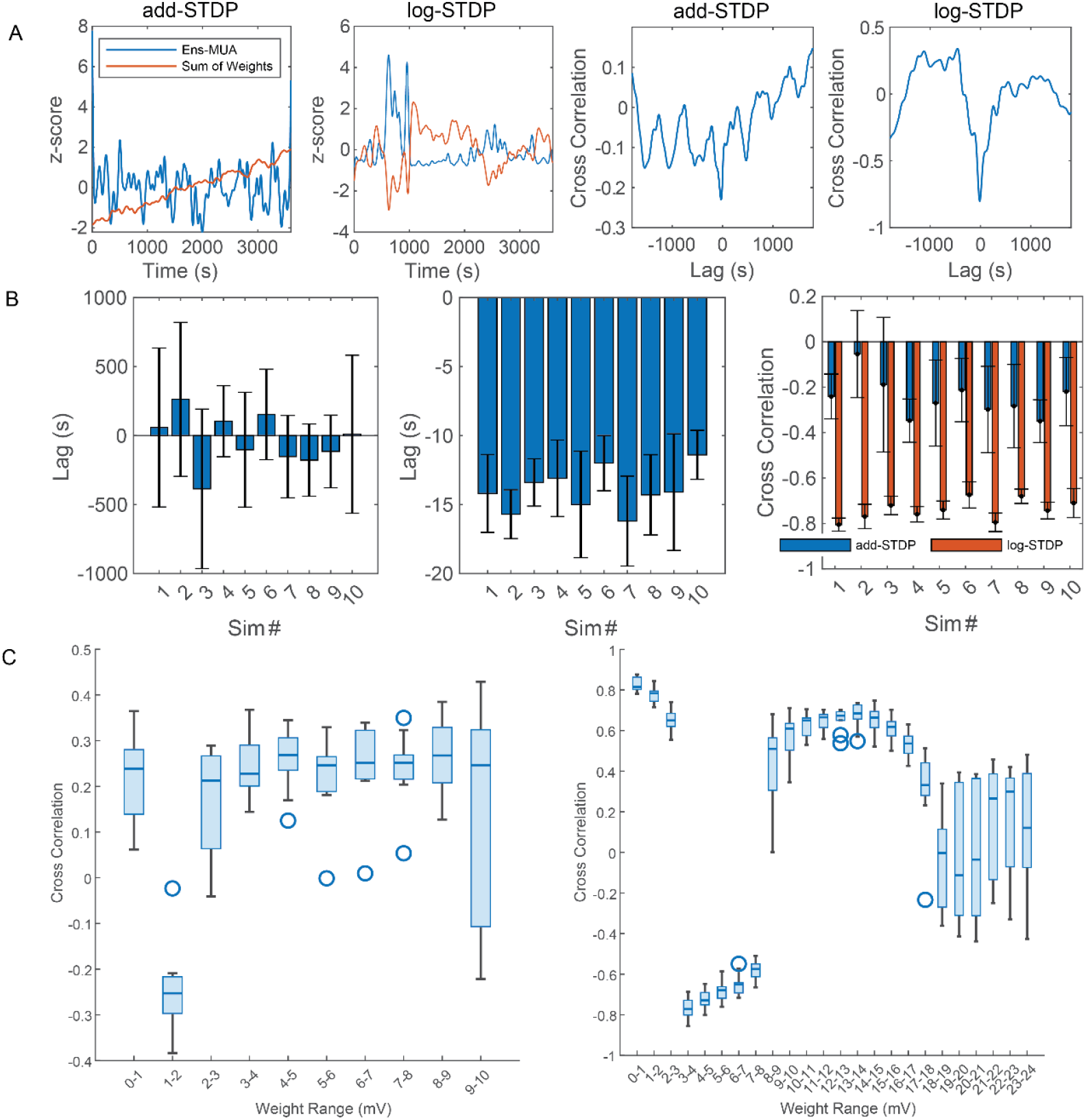
**A.** Examples of smoothed, z-scored traces of ens-MUA and total excitatory weights from CTX simulations (left; middle-left) and there corresponding cross-correlation curves (middle-right; right). Example traces are from one simulation. **B**. Mean ± 1 SD lags corresponding to largest peaks in cross-correlation for add-STDP CTX (left) and log-STDP CTX (middle). Mean ± 1 SD of corresponding peak cross-correlation values for both add-STDP and log-STDP CTX simulations (right). **C**. Cross-correlation values between ens-MUA and the number of connections in specified, 1 mV ranges. Boxes show median and IQR, whiskers show non-outlier maximum and minimum cross correlation values. Outlier threshold was set to 1.5*IQR.

Comparing the total amount of excitatory weight to the ens-MUA does not provide any information regarding how the weight distribution is changing overtime. For instance, when the total amount of excitatory weight decreases does that imply that all connection weights are decreasing in value. To obtain further insights into which weight values were strongly correlated with the CE dynamics, we repeated the same analysis using the number of connections that fall within 1 mV weight ranges instead of the total excitation (see Methods). Most notable from this analysis was the strong correlation of the connections with weights between 0-3 mV and 8-18 mV and the strong anti-correlation for connections with weights from 3-8 mV in the log-STDP CTX simulations (Fig. 5C, right). For add-STDP, there was a weak anti-correlation with the number of weights between 1-2 mV but a weak correlation for all other ranges (Fig. 5C, left). This result indicates that the connections weights in the log-STDP CTX simulations are not undergoing a uniform change in response to changes in the number of active CEs but instead a consistent trend of weights being pushed away from the peak of the distribution.

## 4. Discussion

In this study we investigated the relationship between plasticity and memory by simulating and comparing spiking neural networks with two different STDP rules, add-STDP and log-STDP, and two different network architectures, the cortex and hippocampal CA1. We compared the detection results from anatomically defined memory items PGs, and spike-timing defined memory items CEs. We found the number of PGs detected in simulations using add-STDP were significantly higher than the number found in the log-STDP simulations; however, the opposite was true when comparing the number of detected CEs between the two STDP rules in the CTX simulations; no significant difference was found between the number of CEs in the CA1 simulations. Peak PG sizes were similar across simulations but the maximum PG sizes for the add-STDP CTX simulations were considerably larger. In contrast the sizes of detected CEs were very small for all simulations, with all CEs having fewer than 10 neuron members. The detected CEs in all CTX and CA1 cases had long lifetimes of 3000 seconds, typically. The PGs detected in the add-STDP networks had comparable lifetimes, but the lifetimes of PGs in the log-STDP CTX simulations were shorter and no PGs were detected in the log-STDP CA1 simulations. Lastly, the relationship between the network weights and the number of active CEs was assessed by computing the cross-correlation between the sum of the excitatory connection weights and the number of active CEs each second. The number of active CEs and the total excitatory weight in the log-STDP CTX simulations were strongly anti-correlated with a consistent trend of an increase in the number of active CEs preceding a decrease in the overall amount of available excitation. This strong anti-correlation appears to be driven by the number of weights that fall within the range of 3-8 millivolts in the network. For all other simulations, there were only weak correlations/anti-correlations and very inconsistent lag relationships.

One of the important results from Izhikevich’s (2006) original work was showing that the number of memory items in a network of spiking neurons exceeded the number of neurons in the network. The present study illustrates that this conclusion can change with a different STDP rule. This result has been shown previously (Chrol-Cannon et al., 2012), however, a triphasic STDP rule was used which can still result in a bimodal distribution of the connection weights (Notley & Grüning, 2012). Another factor influencing this conclusion is the choice of definition for the memory item being either anatomically defined (PG) or spike pattern defined (CE). For these simulations, strong connections (>9.5 mV) between neurons are required in order to detect PGs. It follows then that many connections satisfying this requirement imply the existence of many PGs. Furthermore, since PGs require precise temporal coordination of spikes onto a common target it would be reasonable to assume that many PGs mean many precise spike patterns presenting in the spiking activity of the network. The CE results contradict this assumption. Instead, a larger number of strong connection weights in the network results in fewer spike patterns. It is possible that this is due to there being too many strong pre-synaptic connections for every neuron in the network resulting in none of the pre-synaptic neurons being able to reliably stimulate the post-synaptic neuron. Consequently, there is a greater degree of noisy spiking activity in the simulation.

The definition of a PG does not account for the role weaker weights play in developing an association between neurons. In the log-STDP CTX simulations, following an increase in the number of active spike patterns more weights were pushed to the lower end of the distribution (<3 mV) and some were pushed to larger values (>8 mV); this appears to be in line with the idea of stochastic resonance where many weaker connections serve to prepare a postsynaptic neuron by increasing its membrane potential to a point where a single, stronger connection can reliably induce a spike from the postsynaptic cell (Teramae & Fukai, 2014; Teramae et al., 2012). Thus, it is not the convergence of a few spikes onto one cell that is important for producing reliable spike patterns but the convergence of many weak signals with a single stronger signal.

The results for the CA1 simulations suggest a very low capacity for internally generated spike patterns. This, of course, being due to the limited number of recurrent excitatory connections in the network. It may be that the low connectivity of the CA1 is a desirable feature that serves to reduce internally generated noise from impacting the spike patterns projected to the CA1 from other networks.

There are three main limitations to this study to be considered for future work. The first being that the inhibitory connection weights in each of these networks were homogeneous as were the connection delays for inhibitory connections. The connections from inhibitory cells to excitatory cells exhibit their own STDP rule (D’Amour & Froemke, 2015) as do the excitatory-inhibitory connections (Lu et al., 2007). Previous work has illustrated a coordination between the different STDP rules for different connections toward better signal transmission (Kleberg et al., 2014); so, it would be prudent to examine the effect this coordination has on cell ensemble production. The second limitation is the smaller size of these networks. It is possible that the weight distributions are susceptible to network size, for instance, the peak of the lognormal distribution may be considerably lower for larger networks. Finally, the connectivity structure of the network may impact the production and lifetime of CEs in the network. It has been shown previously that the number of PGs in a network of spiking neurons can change depending on network structure (Vertes & Duke, 2010). If the connection delays are taken to be representative of the distance between neurons in the network, then the current method of random assignment can result in situations where there are alternative paths between two neurons through intermediate neurons that are shorter than the direct connection. It is unclear how this impacts the spike patterns present; however, it has been shown that spatiotemporal patterns can be predicted from a weight matrix for a network with time delays corresponding to distance (Budzinski et al., 2023). For future work it would thus be reasonable to enforce distance-based delays.

